# DIffusion-Prepared Phase Imaging (DIPPI): quantifying myelin in crossing fibres

**DOI:** 10.1101/2020.11.10.376657

**Authors:** Michiel Cottaar, Wenchuan Wu, Benjamin Tendler, Zoltan Nagy, Karla Miller, Saad Jbabdi

## Abstract

**Purpose:** Myelin has long been the target of neuroimaging research due to its importance in brain development, plasticity, and disease. However, most available techniques can only provide a voxel-averaged estimate of myelin content. In the human brain, white matter fibre pathways connecting different brain areas and carrying different functions often cross each other in the same voxel. A measure that can differentiate the degree of myelination of crossing fibres would provide a more specific marker of myelination.

**Theory & Methods:** One MRI signal property sensitive to myelin is the phase accumulation, which to date has also been limited to voxel-averaged myelin estimates. We use this sensitivity by measuring the phase accumulation of the signal remaining after diffusion weighting, which we call DIffusion-Prepared Phase Imaging (DIPPI). Including diffusion weighting before estimating the phase accumulation has two distinct advantages for estimating the degree of myelination: (1) it increases the relative contribution of intra-axonal water, whose phase is related linearly to the amount of myelin surrounding the axon (in particular the log *g*-ratio) and (2) it gives directional information, which can be used to distinguish between crossing fibres.

**Results:** Using simulations and phantom data we argue that other sources of phase accumulation (i.e., movement-induced phase shift during the diffusion gradients, eddy currents, and other sources of susceptibility) can be either corrected for or are sufficiently small to still allow the *g*-ratio to be reliably estimated.

**Conclusions:** This new sequence is capable of providing a *g*-ratio estimate per fibre population crossing within a voxel.

## Introduction

Myelin is one of the main constituents of the brain’s white matter^1^ and plays a key role in modulating the speed of action potentials in axons^2,3^. The degree of myelination has been shown to change over a lifetime^4^ with different white matter tracts myelinating at different stages during childhood^5,6^. Activity-dependent changes in myelination have also been demonstrated in adults^7^. The amount of myelin typically decreases during ageing and has been found to be altered in a variety of pathologies^4^, such as leukodystrophies, multiple sclerosis^8^, and schizophrenia^9^. Accordingly, producing accurate *in-vivo* maps of myelin content has been a long-standing goal in brain imaging.

A common metric to quantify the degree of myelination is the *g*-ratio, which is defined as the inner over the outer radii of the myelin sheath^2^. Using multiple MRI modalities one can obtain an estimate of the average voxel-wise *g*-ratio in a voxel *in-*vivo by combining measurements of myelin and axonal volume fractions^10-13^. The axonal volume fraction can be estimated from diffusion MRI, using a multi-compartment fit to the diffusion-weighted signal ^14-18^. A wide variety of different MRI modalities have been proposed to estimate the myelin volume fraction ^19,20^. Most of these rely on directly imaging the myelin water, which can be distinguished from the rest of the water based on its short T_2_* using multi-echo spin-echo sequences^21–23^, its short T_2_* using multi-echo gradient-echo sequences ^24,25^, its short T_1_ using an inversion-recovery sequence ^26^, or based on magnetisation transfer between the myelin macromolecules and water ^27^.

The interpretability of estimating the *g*-ratio from volume fractions is limited, as it only gives an average *g*-ratio per voxel. It is an average across both myelinated and unmyelinated axons ^28^, as the method assumes that all axons have the same *g*-ratio ^11^. It is also an average across fibre populations in voxels where multiple fibres cross each other, which is a common configuration in the human brain ^29,30^. Furthermore, this approach relies on the accuracy of the volume fraction estimates ^31^, which has been questioned for both the axonal volume fractions^32^ and the myelin volume fractions ^13,19,20^. Here we aim to overcome these limitations by proposing a novel sequence, which is directly sensitive to the *g*-ratio (rather than the volume fractions) and allows to distinguish between crossing fibres.

Diffusion-weighting gradients can be used to distinguish between crossing fibres. Diffusion-weighting has previously been combined with all of the myelin-sensitive metrics listed above to obtain tract-specific metrics, namely T_2_ ^33–35^, T_2_*^34,36^, T _1_ ^34,37^, and magnetisation transfer ^38^. Unfortunately, diffusion-weighted gradients take such a long time to build up this sensitivity to fibre orientation that there will be very little signal left associated with the myelin water due to its short T_2_^39^. Rather, after diffusion-weighting, the signal will mainly come from water relatively distant from the myelin, which will reduce the sensitivity of the relaxation and magnetisation transfer properties to myelin.

On the other hand, the off-resonance magnetic field generated by the myelin magnetic susceptibility not only affects the local myelin water, but also has an effect throughout the intra- and extra-axonal spaces in nearby tissue ^40–44^. This provides a means to detect the properties of myelin from more long-lived T_2_ species still visible after diffusion weighting. Hence, we propose a sequence called DIffusion-Prepared Phase Imaging (DIPPI), where we estimate the myelin-induced phase accumulation in the MR signal still visible after diffusion weighting.

In this work we first derive how the phase accumulation measured by DIPPI is related to the *g*-ratio in crossing fibre bundles. We then use simulations and phantom data to show under which conditions we can reliably estimate the myelin-induced phase accumulation and hence the *g*-ratio from DIPPI, despite many potential confounds, namely eddy currents, non-myelin sources of susceptibility, and remaining signal from extra-axonal water after diffusion weighting.

## Theory

### Overview

The DIPPI sequence consists of a standard diffusion-weighted spin echo sequence to which we have added an additional refocusing pulse and readout. The acquisition window of the second readout is offset from the second spin echo by a tuneable delay, which we refer to as the phase accumulation time *t*_phase_ (Figure 1A). The phase difference between these two readouts allows us to estimate the off-resonance frequency of the water still visible after diffusion weighting without being confounded by any phase accumulation during the diffusion weighting.

**Figure 1.**
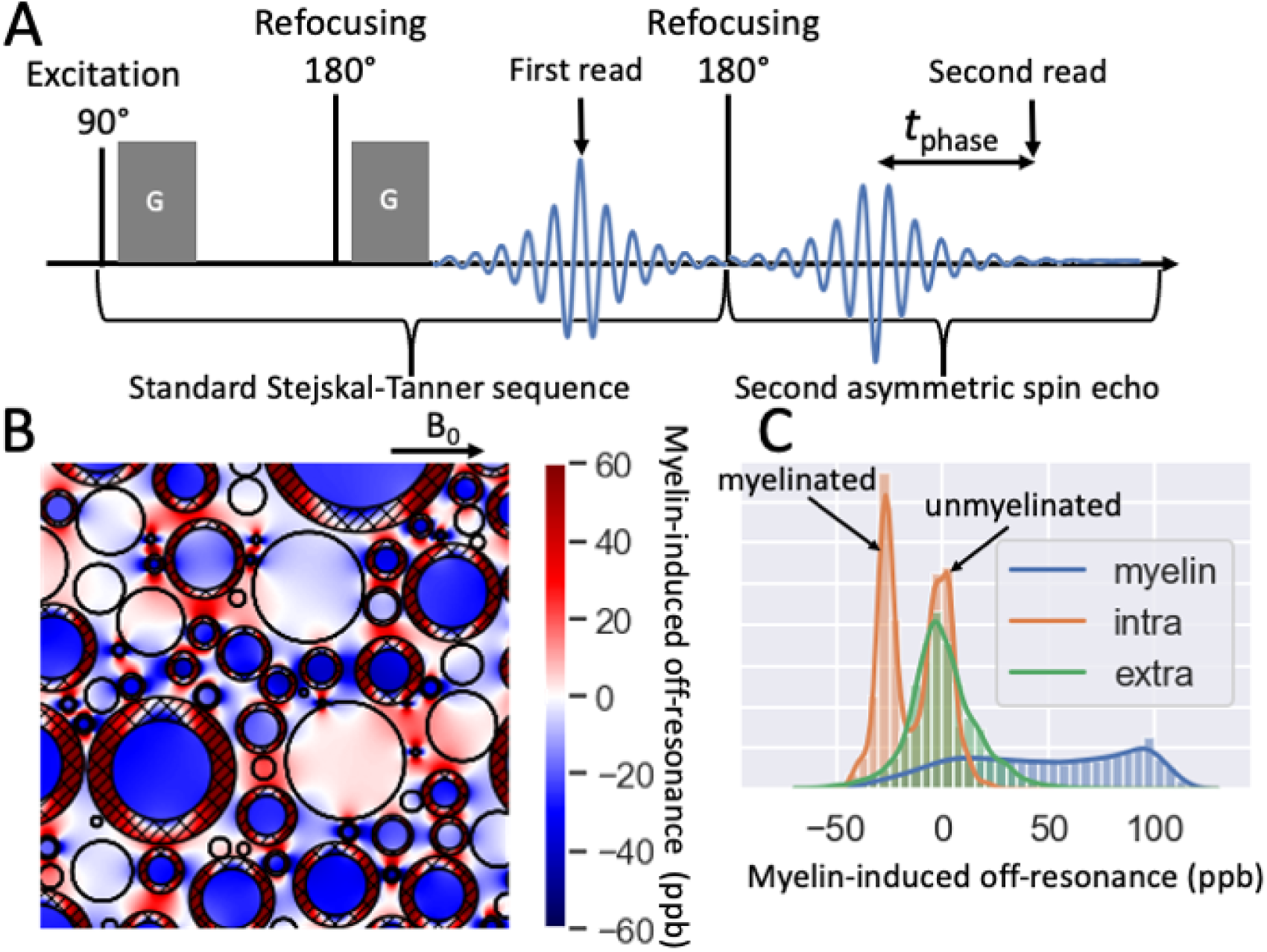
A. Proposed DIPPI sequence to measure the off-resonance frequency of diffusion-weighted water. The sequence consists of a standard Stejskal-Tanner sequence followed by a second EPI readout in an asymmetric spin echo. B. Illustration of white matter with axons as parallel cylinders, some of which are myelinated (myelin sheaths are hashed). Overlaid is the off-resonance field induced by the myelin according to the hollow cylinder model^40^. C. The distribution of the field shown in B in the intra-axonal (orange), extra-axonal (green), and myelin (blue) compartments. After diffusion-weighting the signal will be dominated by the intra-axonal water in axons perpendicular to the diffusion-weighting gradient. For this intra-axonal water, the off-resonance frequency has a bimodal distribution corresponding to the unmyelinated and myelinated axons with the latter having an off-resonance frequency proportional to the log *g*-ratio.

Combining diffusion-weighting with phase imaging provides two advantages to measure the degree of myelination of individual tracts. Firstly, it increases the relative contribution of the intra-axonal water to the final signal, particularly at high b-values ^45^. This has the advantage that while the myelin-induced magnetic field offset has a complicated spatial profile in the extra-axonal and myelin space (Figure 1B,C), it is uniform within the intra-axonal space. For a simplified model of myelinated axons as infinite cylinders, this myelin-induced off-resonance frequency in the intra-axonal space (*ω*_myelin_) is given by^40^:

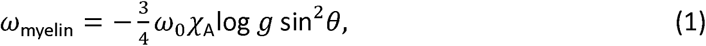

where *ω*_0_ and *χ*_A_ are constants (respectively, the Larmor frequency and the anisotropic component of the myelin susceptibility) and *θ* is the angle between the fibres and the main magnetic field, which we estimate using the magnitude data from DIPPI. The second advantage of using diffusion weighting is that it adds directional information, which allows us to measure the relative degree of myelination (i.e., log *g*-ratio) between crossing fibres rather than a voxel-wide average.

With DIPPI is that we can also exploit the bimodal distribution of the intra-axonal off-resonance frequency (Figure 1C) to fit a two-population model to data acquired with multiple phase accumulation times (*t*_phase_). While for a single *t*_phase_, we can obtain the average log *g*-ratio across both the myelinated and unmyelinated axons, the two-population model allows us to estimate their relative signal fractions axons as well as the average log *g*-ratio of the myelinated axons.

In order to explain the analysis, we split it into three parts. First, we estimate the susceptibility-induced off-resonance frequency of diffusion-weighted water taking into account other sources of phase accumulation (i.e., movement during the diffusion encoding and eddy currents). Then we discuss how to subtract out the off-resonance frequency due to susceptibility sources other than myelin. Finally, we relate the myelin-induced off-resonance frequency to the average log *g*-ratio of crossing fibres.

### Estimating the off-resonance frequency

The DIPPI signal is modulated by both the diffusion-weighting gradients (i.e., the *b*-value and orientation *ĝ*) and the phase accumulation time *t*_phase_. For each set of b-value, gradient orientation, and *t*_phase_, we acquire two images, one during the initial spin echo readout (*S*_SE_) and one during the second asymmetric spin echo readout (*S*_ASE_). In this work we assume that all data have been acquired with a single b-value (in addition to *b* = 0 scans), although the model can be extended to multiple b-values by fitting all parameters independently at each b-value, except for the fibre orientations and degree of myelination.

For a single *t*_phase_ the expected signal across multiple gradient orientations is given by:

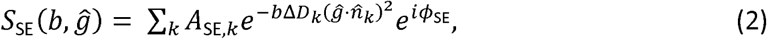

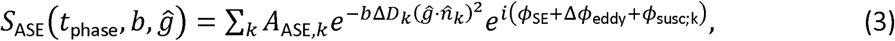

where we sum the signal contributions from multiple crossing fibre populations *k* in an effort to estimate the phase due to the off-resonance frequency associated with each fibre population *ϕ*_susc;k_. The other terms are explained below.

The first part of these equations (i.e., 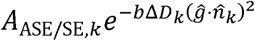) is concerned with the magnitude of the image (Figure 2B). As we are mainly interested in the phase, we fit to the magnitude the simplest model that can distinguish between crossing fibres, namely one where the signal profile for each crossing fibre is given by a Watson distribution with an amplitude *A*_*k*_ and width *ΔD*_*k*_. This is the signal profile expected if the signal for each fibre population can be modelled by an axisymmetric diffusion tensor with eigenvalues *λ*_*‖,k*_ and *λ*_⊥*_,k*_ and volume fraction *f*_*k*_. n that case the amplitude corresponds to 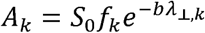 and the width to *ΔD*_*k*_ = *λ*_*‖,k*_ − *λ*_⊥,*k*_.

**Figure 2.**
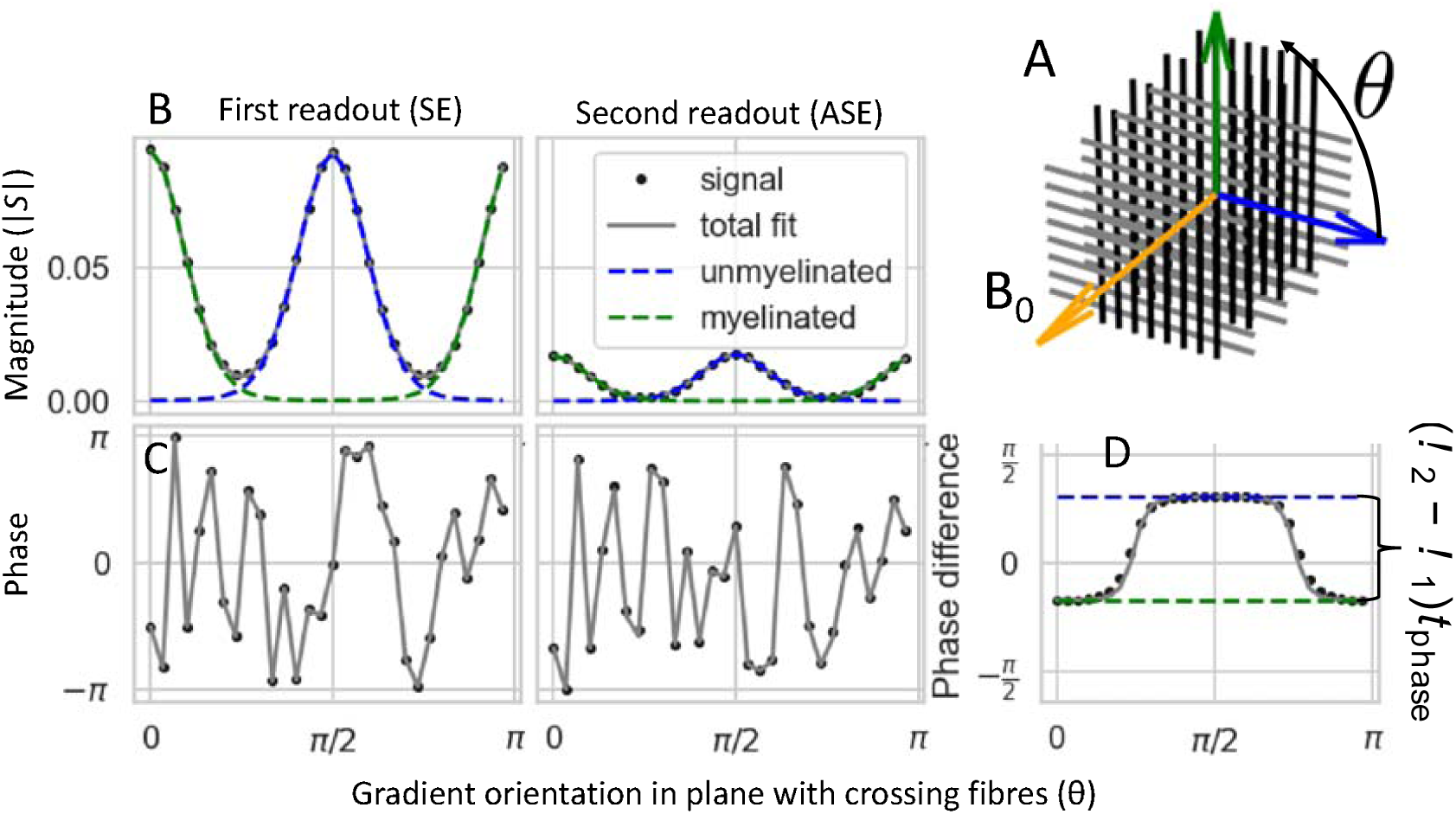
Illustration of the signal estimated from Monte Carlo simulations of two fibre populations (one fully myelinated with a g-ratio of 0.7 and one fully unmyelinated) crossing at right angles and perpendicular to the main magnetic field (A). For ease of illustration we only consider gradient orientations in the plane of the crossing fibres, but the same principle holds for a 3D acquisition. The magnitude is fitted as a sum of 2 Gaussians (Watson distributions in 3D), which have maxima perpendicular to the fibre orientation (B). These Gaussians will have a much lower amplitude in the second readout, but are assumed to have the same width between the readouts. While the phase will be different for each gradient orientation due to movement during the diffusion weighting (C), the phase difference between the two readouts still provides an estimate of the difference in susceptibility-induced off-resonance frequency of the two fibre populations (D).

The width of these Watson distributions (*ΔD*_*k*_) only depends on the diffusion weighting and hence should be the same for both the symmetric and asymmetric spin echoes. The signal amplitudes (*A*_*k*_) on the other hand will decrease over time due to *T*_*2*_ and 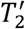 dephasing, which means that we will have a different amplitude for each readout: *A*_SE,*k*_ and *A*_ASE,*k*_. Using multiple phase accumulation times, it is possible to use the dependence of *A*_ASE,*k*_ on *t*_phase_ to estimate both the *T*_2,k_ and 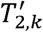 of the diffusion-weighted signal for each fibre population.

The phase accumulation before the first readout will be affected by many factors, such as eddy currents or movement during the diffusion encoding larger than a few tens of micrometres. As such movements are unavoidable in *in-vivo* MRI we simply consider the phase at the first readout to be a random number which has to be estimated independently for each volume (*ϕ*_SE_). Our interest here is in the phase accumulation between the two readouts, which is induced by the off-resonance frequency of any eddy currents (Δ*ϕ*_eddy_) and the brain’s susceptibility (*ϕ*_susc_) (Figure 2C,D).

### Eddy-current induced off-resonance frequency

Eddy currents caused by the strong diffusion gradients will introduce a phase offset that is dependent on the gradient amplitude and orientation. Here, we are interested in the contribution of eddy currents to the phase accumulated between the two readouts

(Δ*ϕ*_eddy_). We model this phase offset using spherical harmonics:

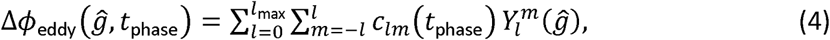

where 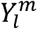 are the spherical harmonic functions mapping the parameters *c*_*lm*_ onto the sphere. Because the eddy currents decay over time following the diffusion-weighting gradients, we cannot simply model these parameters using a linear equation as we will for the susceptibility below.

We can estimate part of the eddy current contributions, because the other contributions to the phase accumulation will be symmetric (i.e., they are identical for a gradient orientation *ĝ* or its inverse − *ĝ*). This means that we can estimate the odd-order spherical harmonics (which are asymmetric), but not the even-order spherical harmonics (which are symmetric and hence degenerate with the susceptibility-induced phase offsets). Fortunately, the dominant component of the eddy current induced phase offset is asymmetric as we will confirm in the Results section.

One exception, where we can estimate part of the even-order components of the eddy-current induced phase offset, is if we acquire a shell with *t*_phase_ = 0 (i.e., both readouts are at their respective spin echoes). For this shell the susceptibility-induced phase offset is zero, so we can attribute any phase accumulated between the two readouts to the eddy currents and hence estimate the even components of *c*_*lm*_*(t*_phase_ = 0). Then, rather than assuming that the even-order components of *c*_lm_*(t*_phase_) are zero we can instead model them by assuming they match *c*_lm_*(t*_phase_ = 0). This corrects for any eddy-current induced phase accumulation between the spin echoes, although it still cannot correct for the evolution of the even components of the spherical harmonics during the phase accumulation time.

### Correcting for the non-myelin susceptibility

The susceptibility-induced off-resonance frequency will not only be influenced by the local myelin (*ω*_myelin_), but also by many other sources of susceptibility (*ω*_bulk_):

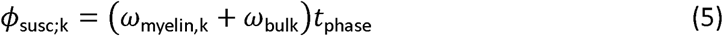

These other sources of susceptibility include both distant sources (e.g., the air-tissue interface) and other local sources of susceptibility (e.g., blood vessels). To resolve between these myelin and non-myelin susceptibility, we make the assumption that any non-myelin source of susceptibility (i.e., *ω*_bulk_) is equal for all crossing fibres. This allows us to estimate the myelin-induced frequency offset difference between crossing fibres (with indices k and k’) as:

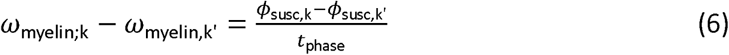

This assumption will be most accurate if the crossing fibres overlap spatially (i.e., they interdigitate). On the other hand, if the crossing fibres are on opposite sides of a voxel, their off-resonance frequency may differ due to any large-scale magnetic field gradients or differences in local susceptibility field (e.g., one fibre population being closer to blood vessels).

Equation 6 only gives the difference in the myelin-induced frequency offset between crossing fibres, which would only allow one to estimate the difference in myelination between crossing fibres. To obtain an absolute estimate of the *g*-ratio for each individual fibre we need additional information. This can be obtained by changing the head orientation, which modulates the relation between the off-resonance frequency *ω*_myelin_ and the *g*-ratio (equation 1). Once the frequency offset (equation 6) has been estimated for multiple head orientations, the individual *g*-ratios can be obtained through linear regression.

### Estimating the *g*-ratio

One additional obstacle to estimating the *g*-ratio is that while there is a simple linear relationship between the myelin-induced off-resonance frequency and the *g*-ratio within each axon (equation 1), each fibre population consists of many axons with potentially varying *g*-ratios. We propose two methods to still obtain a meaningful estimate of the *g*-ratio.

The first method is only valid for *t*_phase_ short enough that the signal phase from the most myelinated axons is still in rough alignment with the signal phase from the least myelinated axons (i.e., the unmyelinated axons with a *g*-ratio of 1 and hence *ω*_myelin_ = 0). In that limit the myelin-induced phase accumulation is determined by the average of the off-resonance frequency in each axon (weighted by its signal contribution) and hence we have:

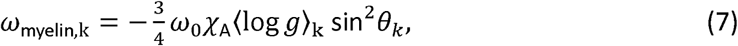

where ⟨log *g*⟩_*k*_ is the signal-weighted average log *g*-ratio of the fibre population k across both myelinated and unmyelinated axons.

For longer *t*_phase_ this simple relation above no longer holds and we need to adopt a two-compartment model: the myelinated and unmyelinated fibres (Figure 1C). For the myelinated fibres we assume that the g-ratios are sufficiently similar that we can characterise this population based solely on their average log *g*-ratio. Hildebrand and Hahn^46^ found a range of g-ratios from 0.6 up to 0.75 in the spinal cord of various mammals. Because this is quite a narrow range compared with the g-ratio of 1 for unmyelinated fibres, we expect the two-compartment model to be adequate for any reasonable *t*_phase_ (at least in healthy tissue). For this two-compartment model we expect a phase evolution of:

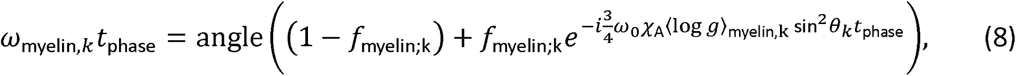

where *f*_myelin;k_ is the relative signal fraction of myelinated versus unmyelinated axons and ⟨log *g*⟩_myelin;k_ is the average log *g*-ratio of just the myelinated axons and “angle” is a function that returns the angle of a complex number. Equation 7 is the first-order Taylor expansion of equation 8 with the average log *g*-ratio across all axons defined as ⟨log *g*⟩_*k*_ - *f*_myelin;k_ ⟨log *g*⟩_myelin;k._

The phase evolution of the signal phase according to equation 8 is shown in Figure 3. At small *t*_phase_ the phase evolution is approximately linear with a slope of *f*_myelin_ *ω*_myelin_, however as myelin phase approaches the phase starts to approach the phase within just the dominant population (i.e., unmyelinated axons for 0 < *f*_myelin_ < 0.5 or myelinated axons for 0.5 < *f*_myelin_ < 1 (Figure 3B). By combining data across multiple *t*_phase_ we can capture this time-dependent non-linear phase evolution to characterise both the fraction of myelinated axons (*f*_myelin_;k) and their average log *g*-ratio (⟨log *g*⟩_myelin;k_) for each crossing fibre population. The evolution of the magnitude also contains information on the myelination (Figure 3C, but in practice this will be very hard to disentangle from other sources of 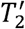 dephasing, which we do not consider here. For this reason, we will constrain the myelination purely on the phase and not the magnitude information. Table 1 summarises which parameters can be estimated for different acquisition schemes.

**Table 1.** Acquisition requirements for the parameters of interest

| Acquisition<br>(all single b-value) | What can be estimated |
| --- | --- |
| Single non-zero $t_{\text{phase}}$ | $\langle \log g_1 \rangle \sin^2 \theta_1 - \langle \log g_2 \rangle \sin^2 \theta_2$ |
| Multiple head orientations | $\langle \log g \rangle_k$ per fibre population |
| Short and long $t_{\text{phase}}$ | $f_{\text{myelin};k}$ and $\langle \log g \rangle_{\text{myelin};k}$ instead of $\langle \log g \rangle_k$ |
| Any above *and* $t_{\text{phase}} = 0$ | Also: $T_{2,k}$, $T'_{2,k}$, $\phi_{\text{eddy, sym}}(t_{\text{phase}} = 0)$ |

**Figure 3.**
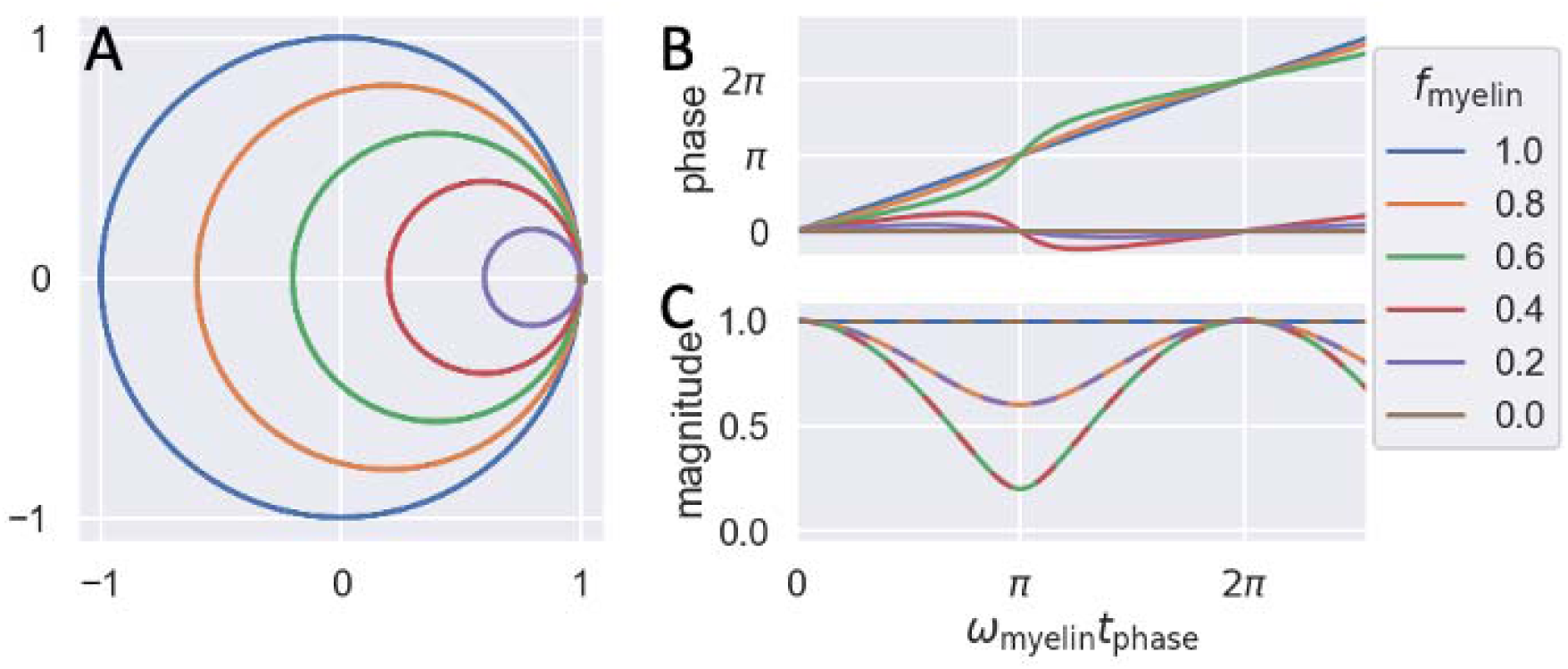
Signal evolution over time for the sum of unmyelinated axonal water (*ω* = 0) and myelinated axonal water (*ω* = _myelin_). Each line shows the evolution for a different signal fraction of myelinated axons (*f*_myelin_; colour coded according to legend on the right). Panel A shows the signal evolution through complex signal space with B and C showing just the phase or magnitude evolution. For only myelinated axons (*f*_myelin_ = 1 in blue) the signal traces a circle in complex space with constant magnitude and linearly increasing phase. As the fraction of unmyelinated axons increases the size of this circle shrinks and importantly it no longer centres on the origin, which leads to non-linear phase and magnitude evolution.

A summary of the model with definitions for all parameter is given in the supplementary materials (S1 with a description of the fitting procedure in S2.

## Methods

### Phantom scan

The DIPPI sequence was implemented on a 7T Siemens scanner. To validate the sequence and to characterise the influence of eddy currents we scanned an isotropic oil phantom. Because the phantom is isotropic, we will attribute any variation in the signal phase between different gradient orientations to eddy currents, which allows direct estimation of their contribution. Three axial slices were acquired using the sequence shown in Figure 1A with the following scan parameters: image resolution 2 mm x 2 mm, slice thickness 2 mm, field of view 192 mm x 192 mm, 6/8 partial Fourier, 10 mm slice gap, echo spacing 0.81 ms, b-value 2 ms/μ^2^m, 60 diffusion directions and their reverse were acquired (i.e., 120 diffusion weighted images in total) and 8 *b* = 0 volumes. The effective echo times for the two readout were 81 and 165 ms, respectively (*t*_phase_ = 30 ms),. After phase unwrapping across gradient orientations (described in the supplementary materials S3), the phase offset observed in the *b*= 0 images was subtracted. Spherical harmonics were then fitted to the phase to estimate the *c*_*lm*_ in our eddy current model (equation 4).

### Reference susceptibility-weighted imaging

To quantify the magnitude of the off-resonance field including all sources of susceptibility, we used publicly available phase imaging data from the QSM reconstruction challenge in Graz^47^. This dataset was acquired from a healthy volunteer using a wave-CAIPI sequence^48^ with an isotropic resolution of 1.05 mm and echo time of 25 ms on a 3T MRI scanner. The provided data has already been phase unwrapped. We convert the phase image to frequency by dividing it by the echo time. Then we compute the magnitude of the local frequency gradient. This gradient gives a rough idea of how different the off-resonance field might be for fibre populations on opposite sides of a voxel.

### Simulations to test extra-axonal contribution

The proposed model assumes that any remaining signal after diffusion-weighting is intra-axonal. To investigate potential biases due to any extra-axonal signal remaining we ran Monte Carlo simulations using Camino’s datasynth^49^ of crossing fibres using the default diffusivity of 2 μm^2^ /ms. Fibres were crossing at 90 degrees (in the x- and y-direction) with both being perpendicular to the main magnetic field (in the z-direction). All axons were modelled as perfect cylinders in the x-direction or y-directions organised in interleaving single-axon thick planes (Figure 2A). The distance between planes was fixed to 1 micrometre. By varying the outer axonal diameter between 0.5 and 0.98 micrometre, we vary the extra-axonal volume fraction from 0.25 to 0.8. Within each plane half of the axons were myelinated (*g* = 0.7), with the other half being unmyelinated. The trajectory of 100,000 simulated spins was output.

The spin evolution over the sequence including the effect of the myelin susceptibility was modelled for a 7T scanner at multiple different b-values. The myelin-induced off-resonance frequency was modelled according to the hollow-fibre model^40^ with myelin susceptibility of *χ*_I_ = −100 ppb (isotropic component) and *χ*_A_ = −100 ppb (anisotropic component). In this model the off-resonance field at every point is evaluated as the contribution of the surrounding axon’s myelin (if any) given by equation 1 and the sum of the dipole-like extra-axonal field of all other axons. The simulated data was fit using the procedure described in supplementary material S2 to estimate the bias due to the signal contribution from extra-axonal water. The confounds of eddy currents, non-myelin contributions to the susceptibility, and measurement noise were not included in these simulations. In addition, the myelin water itself was not explicitly modelled as its contribution is expected to be very small due to its short T_2_ (in fact the border between the intra- and extra-axonal water was infinitely thin and non-permeable in the simulations).

### Simulations to test degeneracy between parameters

Finally, we model and then fit DIPPI data using the model described in the Theory section to investigate any degeneracies between parameter estimates. In these simulations the initial amplitudes and signal widths are set assuming a stick-like diffusion model 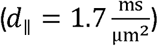, the phase at the first readout (*ϕ*_SE_) is set to a random value between 0 and 2*π* for each scan and the off-resonance frequency due to non-myelin susceptibility (*ω*_other_) is set to a random large value (so that the phase wraps many times between each *t*_phase_). The *l* = 1 components of the eddy currents are computed from *a* + *b t*_phase_, where a and *b* are random numbers drawn from Gaussian distributions *N*(0, *σ* = 1.4 rad) and *N*(0, *σ* = 18 Hz) respectively. We set *T*_2_ = 60 ms^50^ and 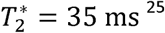 as appropriate for 7T. We consider two crossing fibres at 90 degrees, both of which have 50% of the axons being myelinated with a *g*-ratio of 0.7. Either both fibre populations are perpendicular to the main magnetic field or one is parallel and the other is perpendicular to the main magnetic field. We also consider the case where we have data for both of these fibre configurations with respect to the main magnetic field, which is an extreme case of what can be achieved by scanning with multiple head orientations.

## Results

### Magnitude of myelin-induced off-resonance frequency

To understand the biases induced by phase accumulation due to other sources than myelin susceptibility, we first need to investigate how large the contrast in myelin-induced frequency is expected to be between crossing fibres. Within axons the off-resonance frequency reaches its maximum (*ω*_max_) for fibres perpendicular to the main magnetic field (i.e., sin^2^ *θ* = 1). For a *g*-ratio of 0.7, this maximum off-resonance frequency is about 27 ppb, which corresponds to − 3.4 Hz at 3T and − 8 Hz at 7T (Figure 4A).

**Figure 4.**
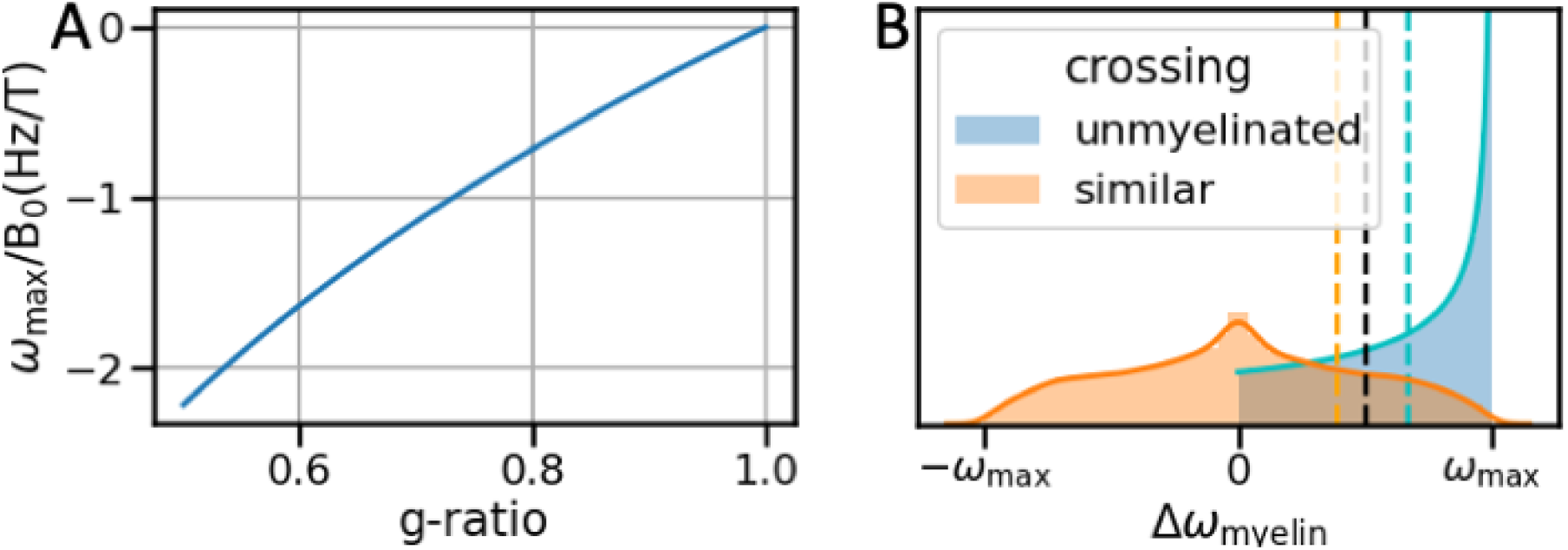
Distribution of the expected difference in off-resonance frequency due to myelin for two crossing fibres. A) maximum off-resonance frequency obtained for fibres perpendicular to the main magnetic field as a function of *g*-ratio. B) distribution of off-resonance frequency difference for two randomly oriented crossing fibres relative to the maximum off-resonance frequency shown in panel A. The distribution is shown for crossing fibres that either have the same *g*-ratio (orange) or for myelinated fibres crossing unmyelinated fibres (blue). The coloured, dashed lines show the mean of the *absolute* values of these distributions at 0.4 *ω*_max_ for similar crossing fibres (orange) and 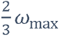 for myelinated crossing unmyelinated fibres (blue). The black dashed line shows the intermediate value (0.5 *ω*_max_) that we adopt as a reference for the typical true effect size for comparison with potential biases in the rest of this work. Only crossings of more than 45 degrees between the two fibre populations are considered.

We compute the myelin-induced off-resonance frequency for two randomly oriented fibre populations (i.e., randomly oriented with respect to each other and the main magnetic field, excluding any fibre crossings with angles less than 45 degrees). Figure 4B shows the distributions of the difference in this myelin-induced off-resonance frequency with respect to this maximum frequency offset for two extreme cases: (1) myelinated axons crossing unmyelinated ones or (2) two identically myelinated populations crossing each other. When myelinated fibres cross unmyelinated fibres, the frequency offset between the two will be fully determined by the angle of the myelinated fibres with the main magnetic field. The offset reaches its maximum of *ω*_max_ when the myelinated fibres are perpendicular to the main magnetic field and minimum of 0 when parallel to the main magnetic field with a mean offset of 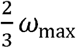 (blue in Figure 4B). For two crossing fibres with identical levels of myelination, the frequency offset will be typically lower with an offset of 0 when the fibres have the same angle to the main magnetic field, while the maximum offset *ω*_max_ is only reached when one fibre is parallel and the other is perpendicular to the main magnetic field. For randomly oriented fibres the mean offset is approximatel myelinated crossing unmyelinated y 0.4 *ω*_max_ (orange in Figure 4B).

This analysis suggests that we can expect the typical difference in myelin-induced frequency offset between crossing fibres to be on the order of half of *ω*_max_, which for a typical *g*-ratio of 0.7 is about 1.7 Hz at 3T and 4 Hz at 7T. Hence, any other uncorrected differences in phase accumulation between the crossing fibres need to be lower than this level to allow reliable measurement of the *g*-ratio.

### Bias due to eddy currents

To investigate the potential bias due to eddy-current induced phase accumulation, we compare the expected myelin-induced phase offsets with the phase offsets induced by eddy currents found in an isotropic phantom on a 7T scanner. The eddy-current induced phase offset is dominated by the *l* = 1 components of the spherical harmonics (Figure 5A). Fortunately, these and the other odd-order components can be estimated even when scanning an anisotropic medium like the brain’s white matter. The even components of the spherical harmonics (for *l* ≠ 0) are more problematic as they are degenerate with the myelin-induced frequency offset between crossing fibres. Fortunately, the power in the even components is typically much smaller than in the odd components (Figure 5B). The only exception is the *l* = 2 component. For comparison, the vertical line shows the magnitude of the expected myelin-induced offset, which we estimated above to be 4 Hz at 7T, which after 30 ms comes to 0.75 rad. For most of the phantom even the l = 2 component is ∼5 times smaller than the myelin-induced offset, which suggests it would not cause a major bias. However, at the edge of the phantom it becomes comparable. This large *l* = 2 component at the edge of the phantom accumulates mostly between the two spin echoes (Figure 5C), rather than between the second spin echo and the readout (Figure 5D). This suggests that it might be better to approximate the *l* = 2 component in a separate scan with t_phase_ = 0 rather than ignoring it altogether.

**Figure 5.**
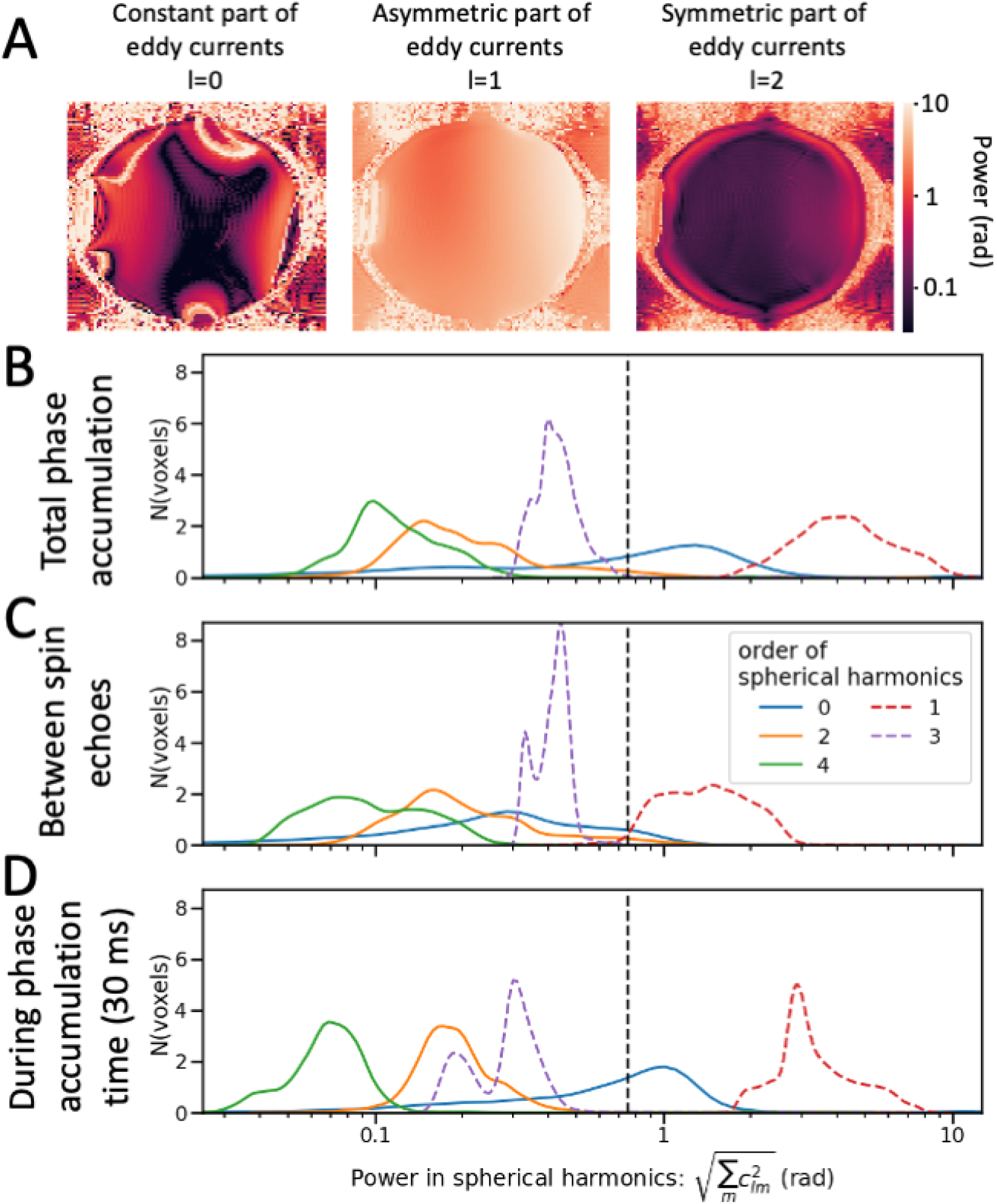
Phase accumulation between the two readouts due to non-zero eddy currents measured in an isotropic oil phantom at *b* = 2 ms/μm^2^ for a phase accumulation time t_phase_ of 30 ms on a 7T Siemens scanner. The phase accumulation measured at b=0 is subtracted out (to subtract out the susceptibility field) and then spherical harmonics are fitted to the eddy-current induced field offset. A. Map of the power in these spherical harmonics. B. Histogram of the maps shown in A. Dashed lines show the odd components (that can be corrected for); solid lines show the even components (that cannot be corrected for). C. Phase accumulation between the first and second spin echoes (measured using t_phase_ of 0 ms). D. Phase accumulation in the 30 ms between the second spin echo and the readout. The dashed vertical line shows the expected approximate magnitude of the myelin-induced phase accumulation expected in the brain.

Unexpectedly, despite subtracting out the phase offset due to the B0 field (estimated using the *b* = 0 scans) the *l* = 0 component is still substantial (Figure 5B). This means that after averaging out all gradient orientations there is a net phase offset on the order of 0.5-1 rad between the *b* = 0 and *b* = 2 ms/μm^2^ scans. The origin of this component is unclear, however we note that it is swamped by the size of the B0 field (discussed below) and will not cause a bias in the frequency offset measured between different fibre orientations.

Figure 6 illustrates the result of the correction of the eddy-induced phase offset between two nearly orthogonal fibre orientations. Without correcting for eddy currents there is a substantial phase offset, which would hide any myelin-induced phase offsets (Figure 6A). Subtracting out the odd-order spherical harmonics gets rid of most of the eddy-current induced phase (Figure 6B). The remaining phase deviations, particularly at the edge of the phantom, are due to the large size of the *l* = 2 spherical harmonic components (Figure 5A) and can be further reduced by subtracting out the phase difference accumulated at *t* _phase_ = 0 (Figure 6C). The histogram of the resulting phase offset does not nicely centre at zero. Although this indicates that a non-zero bias in the phase offset remains, it is small compared with the expected size of the myelin-induced phase offset (indicated by the dashed vertical lines).

**Figure 6.**
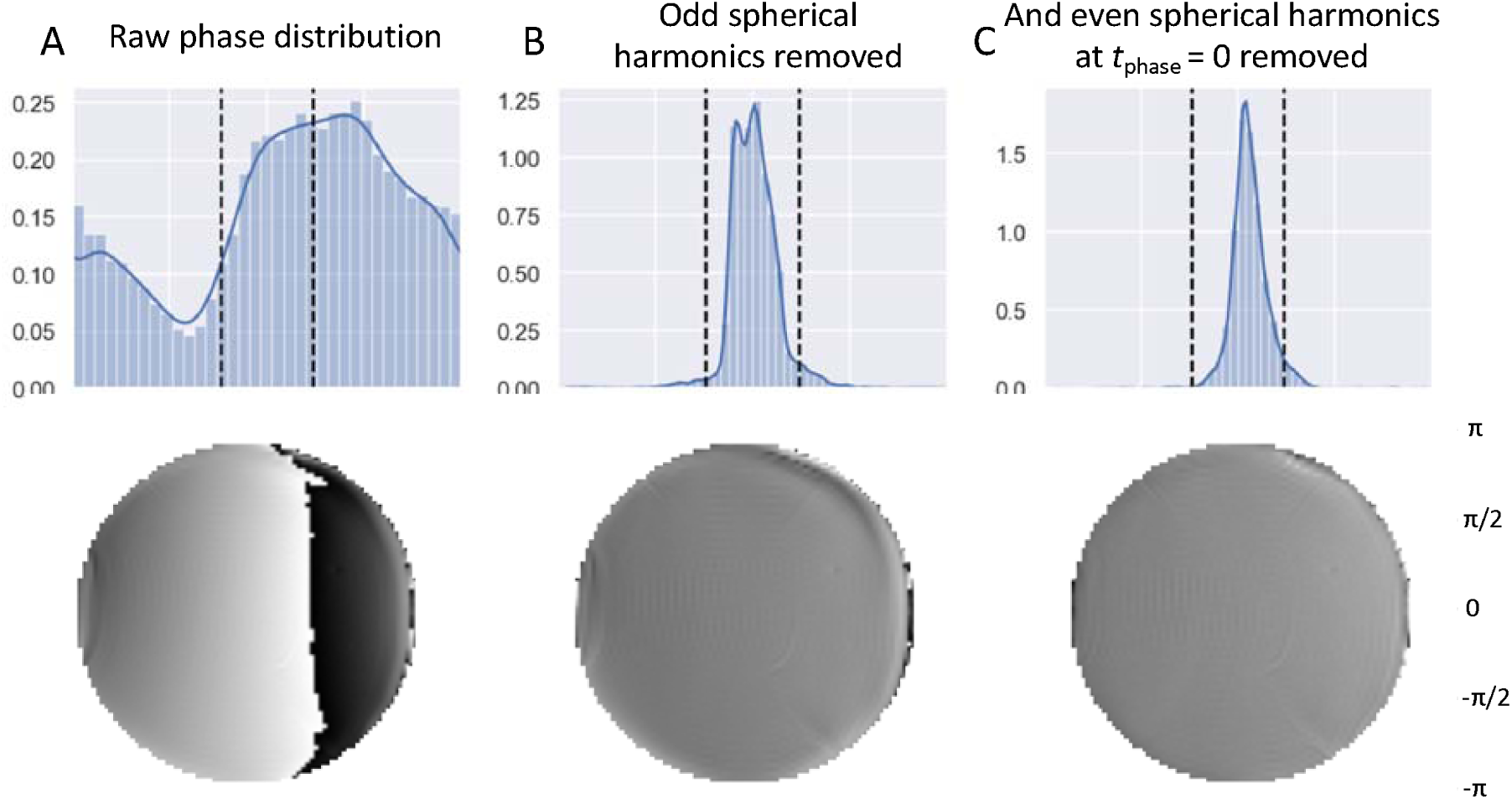
Distribution of difference in the eddy-current induced phase offset between two roughly orthogonal gradient orientations without correction (A), after correcting only the odd-order spherical harmonics (B) or after also subtracting the even-order spherical harmonics estimated at *t*_phase_ == 0 (C). Top panels show the histogram across all three slices. Bottom panels show the phase map for the centre slice. The vertical dashed lines in the top panel show the expected approximate magnitude of the myelin-induced phase offset. In all panels this figure shows the phase difference at *t*_phase_ = 30 ms for an isotropic phantom in a 7T scanner.

### Bias due to bulk susceptibility

The large-scale background off-resonance frequency field is much larger than the expected myelin-induced frequency offset (Figure 7A). This field can bias the *g*-ratio estimate If crossing fibres do not interdigitate, but are actually on opposite sides of the same voxel. In such a case, they may have different contributions from the large-scale off-resonance field. To estimate the size of this effect we computed the spatial gradient of the off-resonance frequency field (Figure 7B). For most of the brain, the gradient of this field is so small that even if the crossing fibres were on the opposite side of a voxel (i.e., about 0.5 mm apart), the resulting frequency offset would still be ∼5 times smaller than that expected from myelin. However, close to the major arteries or the air-brain interface (e.g., the orbitofrontal regions), the gradient of the off-resonance frequency becomes large enough to significantly bias the estimated *g*-ratios in the case that the crossing fibres are not interdigitated.

**Figure 7.**
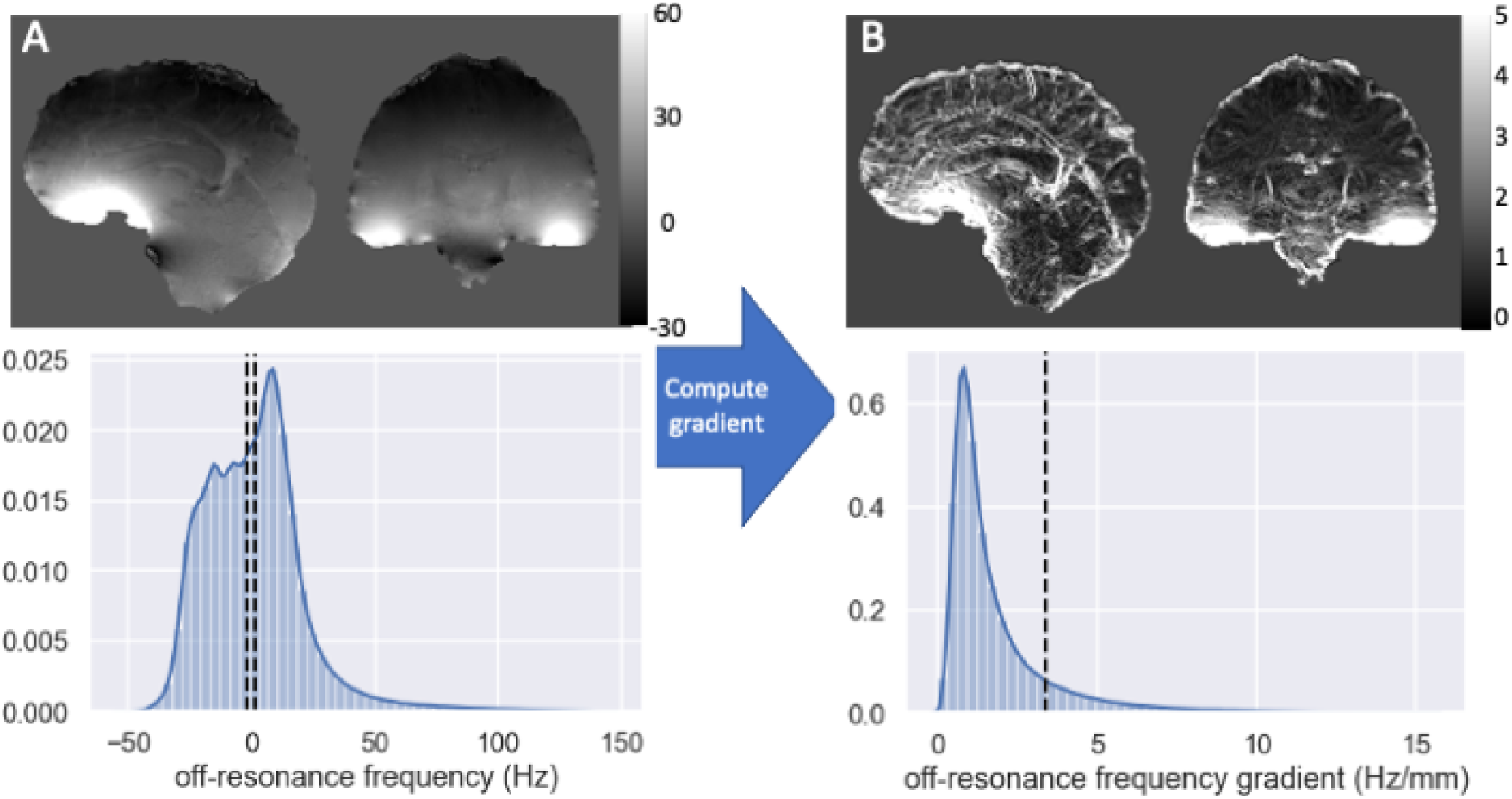
A) Off-resonance frequency distribution estimated for a healthy volunteer at TE=25ms on a 3T scanner (after phase unwrapping) with the dashed vertical lines showing the expected approximate magnitude of the myelin-induced off-resonance frequency difference between crossing fibres. B) Distribution of the gradients in the off-resonance field. The dashed vertical line indicates how large the gradient would have to be to cause an effect as big as expected from myelin if the crossing fibre populations were half a millimetre apart.

### Bias due to extra-axonal water signal

Finally, bias in the estimated parameters can also come from the remaining contribution of the extra-axonal water even after diffusion weighting. For reasonable b-values (∼ 3 ms/μm^2^) we find that the actual off-resonance frequency is ∼10-20% smaller than expected for pure intra-axonal water (Figure 8A), which would lead to a similar underestimation in the log *g*. This underestimation is caused by the ∼15% extra-axonal signal contribution remaining at *b* = 3 ms/μm^2^ (Figure 8B) modulated by the average off-resonance frequency of the extra-axonal water (Figure 8A). Interestingly, in the simulations the 15% extra-axonal signal contribution was consistent across a wide variety of different axonal densities (colour scales). The fibre packing configuration will affect both the average extra-axonal off-resonance frequency^41,51^ and how fast the extra-axonal signal decays with b-value. The simulations here use an unrealistic fibre configuration of perfectly straight cylinders crossing each other at right angles in a perfect grid, which means that the bias found here is a only rough estimate of the bias size expected in real tissue.

**Figure 8.**
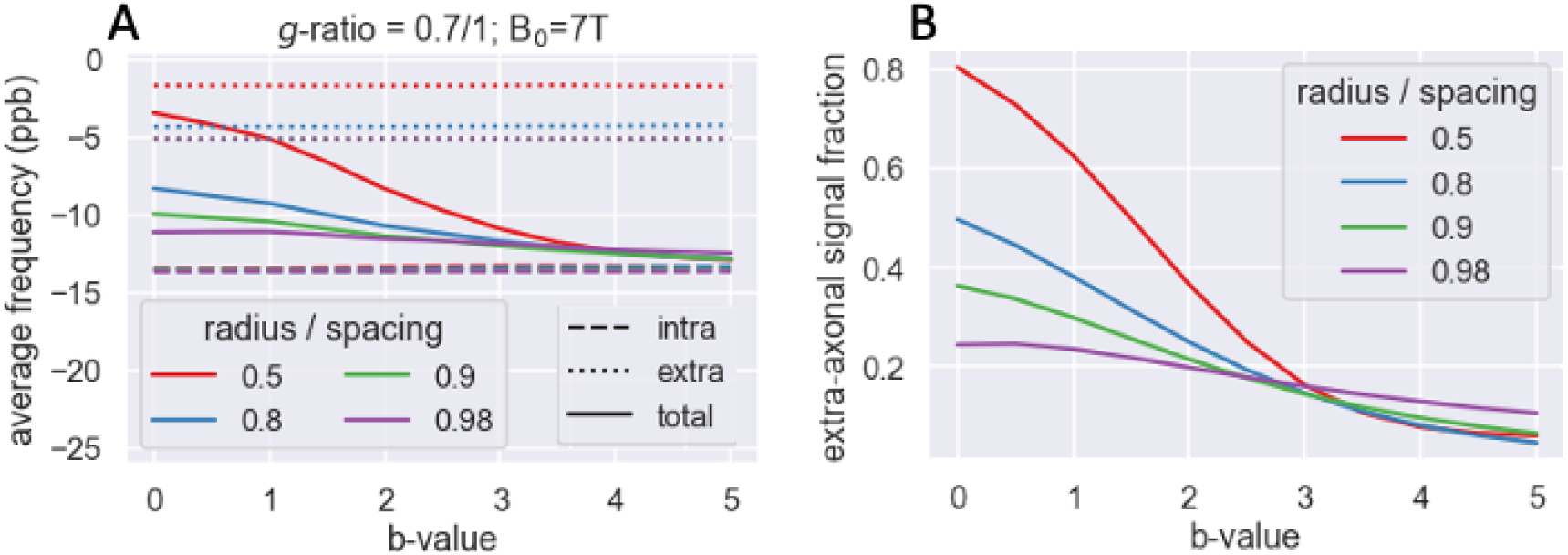
Bias on the off-resonance frequency due to extra-axonal water. A) Average myelin-induced off-resonance frequency in Camino Monte Carlo simulations of infinitely long crossing cylinders (half are unmyelinated and half have *g*=0.7) with different spacings (see colour legend). As the b-value increases the off-resonance frequency of the total signal (solid line) approaches that of the intra-axonal water (dashed), although some bias to the extra-axonal frequency (dotted) remains. B) This approach is caused by the decrease in the extra-axonal signal fraction with b-value in these Monte Carlo simulations.

### Degeneracies between fitted parameters

While the eddy currents, gradients in the non-myelin susceptibility, and extra-axonal water all might bias the estimated *g*-ratios as discussed above, a more fundamental limitation arises because we can only estimate the difference in the myelin-induced frequency between crossing fibres. In case of data only acquired with a single head orientation and single *t*_phase_ we can only estimate a weighted difference in ⟨log *g*⟩ between two crossing fibres (⟨log *g*_l_⟩ sin^2^ *θ*_1_ − ⟨log *g*_2_⟩ sin^2^*θ*_2_). If both fibres have the same angle with the main magnetic field (i.e., *θ*_1_ = *θ*_2_), this implies we can estimate the difference in ⟨log *g*⟩ between the crossing fibres, not what the ⟨log *g*_1_⟩ and ⟨log *g*_2_⟩ actually are. This case is illustrated in Figure 9A by the distributions of blue dots, which all have a very similar ⟨log g_1_⟩ − ⟨log *g*_2_⟩, even while the individual estimates of ⟨log *g*⟩ are unconstrained. On the other hand, if fibres have different angles with the main magnetic field we are less sensitive to the ⟨log *g*⟩ that is more parallel to the main magnetic field, which changes the slope of the degeneracy (i.e., the line along which the points lie in Figure 9A). The most extreme case of this is when one fibre population is parallel to the main magnetic field (e.g., sin *θ*_2_ = 0), in which case we are completely insensitive to the myelination of that population (i.e., ⟨log *g*_2_⟩), but can estimate the ⟨log *g*⟩ of the other population (orange in Figure 9A). By combining information across multiple head orientations, we can constrain the *g*-ratios of the crossing fibres as the intersection between the different degenerate solutions (green in Figure 9A).

**Figure 9.**
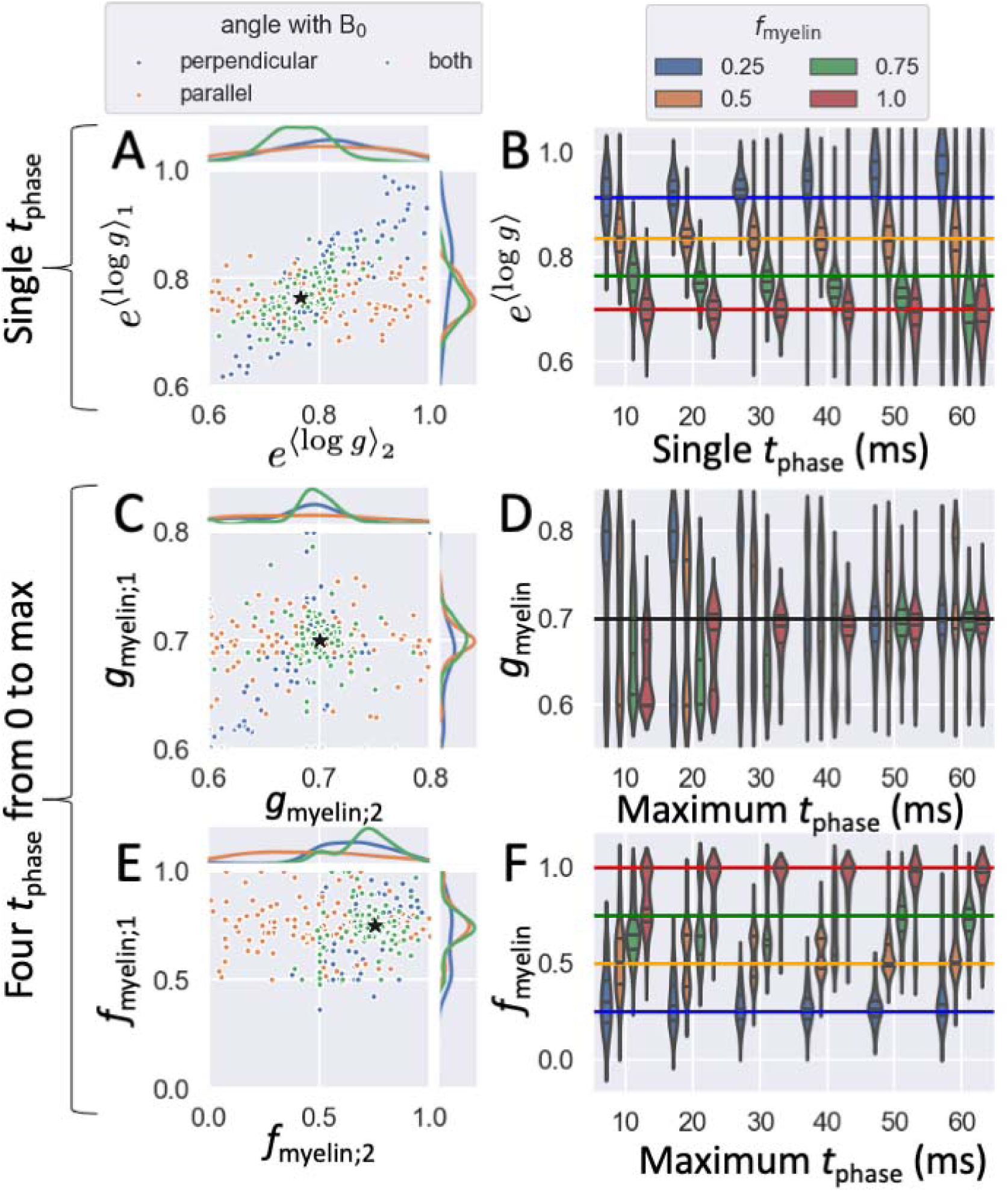
Results for fitting the model with various head orientations (A,C,E) or different phase accumulation times *t*_phase_ (B,D,F). We either consider data with a single *t*_phase_ (20 ms in A), where we estimate the average log g-ratio across both myelinated and unmyelinated fibres (A,B), or data with four different *t*_phase_ uniformly distributed from 0 to the maximum *t*_phase_ (60 ms in C and E) for which we estimate the average log g-ratio of the myelinated axons (*g*_myelin_) and the fraction of myelinated axons (*f*_myelin_). In the left column (A,C,E) each dot represents the estimated value for one of 100 different noise iterations for the case where both fibres are perpendicular to the main magnetic field (blue), one of the fibres is parallel and the other is perpendicular to the main magnetic field (orange), or we have multiple head orientations combining the information from the first two (green). The ground truth value is given by the black star. The right column (B,D,F) shows the distribution of these individual estimates as a function of the maximum *t*_phase_ for various values of the fraction of myelinated axons (*f*_myelin_) for the case of multiple head orientations (green in left column).

Figure 9A considers the case where we aim to estimate the average log g across both myelinated and unmyelinated axons (equation 7) using a single tphase. However, as this *t*_phase_ becomes too long the signal from the unmyelinated and myelinated axons become out of phase with each other and the resulting phase approaches that of the dominant component (Figure 3B). As a result, the estimated average log *g* will then match that of the dominant component (i.e., the myelinated axons if *f*_myelin_ > 0.5 or the unmyelinated axons if *f*_myelin_ < 0.5) (Figure 9B). When we have data across multiple *t*_phase_ we can exploit this behaviour to estimate both the average log g across both components and the log g of the dominant component, which allows the estimation of both the fraction and g-ratio of myelinated axons (Figure 9D,F). In Figure 9D,F we see that this technique works best if the signal is dominated by myelinated axons (i.e., large *f*_myelin_), although as long as the *t*_phase_ is long enough the *f*_myelin_ and to a lesser extent the *g*_myelin_ can be estimated even when the unmyelinated compartment dominates (*f*_myelin_ = 0.25, blue violins plots).

The non-linear time evolution of the myelin-induced phase offset can also be exploited to distinguish it from the non-myelin susceptibility. This leads to reasonable fits to the fraction and g-ratio of myelinated axons even if data was only acquired with a single head orientation (blue in Figure 9C,E). With multiple head orientations, these estimates are still substantially improved (green in Figure 9C,E).

## Discussion

We propose a sequence, DIPPI, to estimate the g-ratio of axons within the white matter by measuring the off-resonance frequency of the water remaining visible after diffusion weighting. After diffusion-weighting the signal is dominated by intra-axonal water in axons that run perpendicular to the diffusion gradient orientation. We exploit the linear relationship between the log g-ratio and the myelin-induced frequency offset in this intra-axonal water (equation 1) to estimate the g-ratio after correcting for several other sources of off-resonance frequency. DIPPI allows one to go beyond the voxel-wise average estimates of g-ratio to get an estimate of the average log g-ratio in every fibre population. For a single, short *t*_phase_ this log *g*-ratio will be averaged across both myelinated and unmyelinated axons, but by varying *t*_phase_ we can separate the myelinated and unmyelinated axons to estimate the volume fraction and g-ratio of the myelinated axons in each crossing fibre population.

Many other sources can affect the amount of phase accumulated in the diffusion weighted signal besides the myelin susceptibility. In DIPPI we measure the phase accumulation between the two readouts after diffusion weighting, which will be unaffected by any phase accumulated during the diffusion weighting (Figure 2). However, phase accumulation between these readouts will still be affected by remaining extra-axonal signal, eddy currents, and non-myelin sources of susceptibility.

In Monte Carlo simulations with greatly simplified geometries we found that the remaining extra-axonal water signal at *b* = 3 μm^2^/ms is around 15% (Figure 8B). To first order, this extra-axonal water has a similar myelin-induced frequency as the water within unmyelinated axons, which suggests that this would lead to an overestimation of the fraction of unmyelinated axons (and a corresponding bias in the average log *g*-ratio; Figure 8A).

The phase accumulated due to eddy currents can be mostly corrected for by modelling them as depending linearly on the gradient orientations. However, not all higher order terms can be so easily corrected. DIPPI data in an isotropic phantom (7T scanner with *b* = 2 μm^2^ /ms) suggests these higher-order terms might bias our *g*-ratio estimates up to about 20% (Figure 6).

The off-resonance field generated by any non-myelin sources of susceptibility is generally much larger than that generated by myelin (Figure 7A). As long as the crossing fibres interdigitate, we can assume that the non-myelin susceptibility contributes equally to their non-resonance fields, which allows us to estimate the difference in myelin-induced susceptibility between the crossing fibres. If the fibres do not interdigitate, but are instead 0.5 mm apart, this assumption could lead to a substantial bias especially close to the air-brain tissue boundary and major arteries (Figure 7B).

Even if the fibres interdigitate, the large size of the non-myelin induced field still means that we can only estimate the difference in myelination between crossing fibres. For data with multiple head orientations we can get around this limitation to get an estimate of the myelination for each crossing fibre population (green in Figure 9A,C,E). However, this technique will not work in single-fibre regions. There are alternative approaches that do not require multiple head orientations. Background field removal might be sufficient to remove *ω*_bulk_ under the assumption that the local susceptibility is dominated by myelin^52–54^ Alternatively, the curvature of white matter tracts naturally varies the angle between the fibre orientation and the main magnetic field, which we can exploit under the assumption that the fibre myelination is constant along the tract. The reliability of these various approaches will be investigated in future work.

When fitting the two-pool model to estimate both the fraction and *g*-ratio of the myelinated axons, additional sources of bias might occur. These estimates rely on the time-dependence of the off-resonance frequency due to the difference in off-resonance frequency between myelinated and unmyelinated axons (Figure 3). Hence, the estimates will be biased by any other sources of time dependence in the off-resonance frequency, which could arise by having multiple compartments with different *T*_2_ and off-resonance frequency. However, we are unaware of any evidence for such time-dependence in the off-resonance frequency at these long echo times.

Finally, we note that there are substantial uncertainties in our estimates of the anisotropic component of the myelin susceptibility (*χ*_*A*_)^43^ limiting the accuracy of the resulting *g*-ratio. A reliable estimate of this constant is crucial to accurately map the *g*-ratio to the intra-axonal myelin-induced frequency offset (equation 1).

The combination of theory, simulations, and phantom data presented here suggests that DIPPI would be able to obtain a reliable measure of the *g*-ratio in crossing fibres. We plan to further explore this using both *in-vivo* and *ex-vivo* data in future work.

## Acknowledgements

MC and SJ are funded by a Wellcome Collaborative Award (215573/Z/19/Z). KM and BT are funded by the Wellcome Trust (202788/Z/16/Z). WW is funded by the Royal Academy of Engineering (RF201819/18/92). The Wellcome Centre for Integrative Neuroimaging is supported by core funding from the Wellcome Trust (203139/Z/16/Z). Human SWI data was used from the QSM reconstruction challenge^46^ with permission from Ferdinand Schweser. We thank Johanna Vannesjo and Jesper Andersson for insightful discussions on the modelling of eddy currents.

## Supplementary Materials

### S1: Model summary

For clarity we provide here a summary of the full model fitted to the complex signal of the first spin echo (SE) readout and the second asymmetric spin echo (ASE) readout:

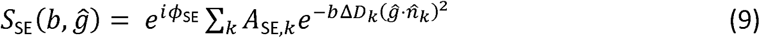

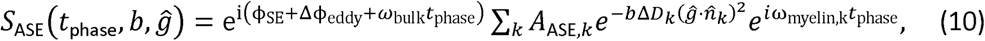

where we sum over multiple crossing fibre populations *k*.

The parameters in this equation are:

- Acquisition parameters
  - *b*: quantifies the sensitivity to the diffusivity
  - *ĝ*: orientation of the diffusion-weighted gradient
  - *t*_phase_ : phase accumulation time between the second spin echo and the centre of the second readout
- Free parameters fitting the signal magnitude
  - *A*_SE_,k: signal amplitude perpendicular to the fibre orientation at the first readout
  - *ΔD*k: signal width corresponding to parallel minus perpendicular apparent diffusivity
  - 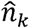: average fibre orientation
  - *A*_ASE_,_*k*_: signal amplitude perpendicular to the fibre orientation at the second (asymmetric) readout, which is fitted independently for each *t*_phase_. With multiple *t*_phase_ can be used to estimate *T*_2;*k*_ and 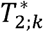.
- Free parameters fitting the signal phase
  - *ϕ*_SE_: any phase accumulated before the first readout. This is estimated independently for each volume.
  - *Δϕ*_eddy_ : eddy-current induced phase offset between the two readouts, which is modelled as a function of gradient orientation using spherical harmonics (equation 4). Odd and even components of the spherical harmonics are treated differently:
    ▪ Odd components of the spherical harmonics are estimated independently for each *t*_phase_
    ▪ Even components are either assumed to be 0 or if only data with *t*_phase_ > 0 is acquired or constant across all *t*_phase_ if data with *t*_phase_ = 0 is also acquired, effectively setting them to the value estimated at *t*_phase_ = 0. Any more realistic model that allows for variability with *t*_phase_ will lead to degeneracies with the estimated *g*-ratio
  - *ω*_bulk_: off-resonance frequency due to non-myelin sources. Assumed to be a constant in each voxel (i.e., does not depend on fibre population or any of the acquisition parameters)
  - *ω*_myelin;k_: myelin-induced frequency offset. If only short *t*_phase_ are acquired, can be related to the average log *g*-ratio (⟨log *g*⟩_k_) through equation 7. Alternatively, a two-compartment fit (equation 8) can be applied if multiple including long *t*_phase_ were acquired. The latter estimates both the signal fraction (*f*_myelin;*k*_) and log *g*-ratio (⟨*log g*⟩_myelin;*k*_) of the myelinated axons.

### S2: Parameter fitting

Parameter estimation is complicated by the phase wrapping inherent in fitting complex MRI data. The signal is identical for a phase of *ϕ* and *ϕ*+ 2 *πn* for any integer *n*, which leads to many unphysical, local minima when fitting the phase. While phase unwrapping could potentially deal with this by estimating *n* for each gradient orientation relative to the others, such unwrapping is complicated by the low SNR inherent in diffusion MRI for gradient orientations aligned with the dominant fibre orientation. Instead, we propose to deal with the phase wraps through a careful initialisation of the parameters and a multi-step fitting procedure. At each step the results of the previous fit are used to initialise the new fit

1. In actual DIPPI data the number of fibres and their orientations can be estimated from any model allowing for crossing fibres such as ball & stick^29^ or constrained spherical deconvolution^56^. In this work, we only simulate data with two crossing fibres and then fit it assuming two crossing fibres. Fibre orientations are initialised in this work to the first two eigenvectors of a diffusion tensor fit.
2. In the initial part of the fitting we only fit the signal magnitude. First, we just fit the amplitudes (*A*_SE_, and *A*_ASE,*k*_ or *T*_2;*k*_ and 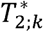), then we fit the amplitudes and width, and finally the amplitude, width and orientation (i.e., the full Watson distribution).
3. Once we have a decent fit for the signal magnitude, we can estimate the parameters influencing the signal phase:
  a. The phase offsets induced during the diffusion encoding (*ϕ*_SE_) are initialized by the phase measured during the first readout. While such direct phase estimates can be very noisy for gradient orientations with very low SNR (≲1), any gradient orientations with such low SNR at the first readout will be so dominated by noise at the second readout that they do not contribute significantly to the final fit.
  b. The *l* = 1 spherical harmonic components of the eddy currents across gradient orientations are then estimated for each *t*_phase_ through the following algorithm
    i. We compute a convex hull containing all the gradient orientations using the Quickhull algorithm^57^ from www.quickhull.org. Any gradient orientations connected in this hull are considered neighbours.
    ii. For each pair of neighbouring gradient orientations the phase difference is computed (and mapped between − *π* and *π* by adding or subtracting 2*π*).
    iii. The linear phase gradient in the x-direction *G*_*x*_ is then estimated by solving: 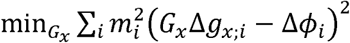, where the sum is over each pair of neighbours *i*, Δ*ϕ*_*i*_ is the phase difference computed in step 2, *m*_*i*_ is the average magnitude of the signal for the neighbouring gradient orientations, and Δ*g*_*x;i*_ is the difference in the x-component of the gradient between the neighbours. The same equation is solved for the y- and z-components. The phase gradient estimates are multiplied by 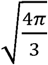 to get the first-order spherical harmonic components.
  c. The non-myelin off-resonance frequency (ω_bulk_) and *g*-ratio parameters are randomly initialised. ω_bulk_ is drawn from a uniform distribution between −300 and 300 Hz, which is a substantially larger range than seen in susceptibility-weighted MRI. For a single-population model the ⟨log g⟩_*k*_ is initialised randomly between log 0.6 and log 1 (= 0). For the two-population model *f*_myelin;*k*_ is initialised randomly between 0 and 1 and ⟨log g⟩_*k*_ between log 0.6 and log 0.8.
  d. We then fit in order just the non-myelin off-resonance frequency (ω_other_), the non-myelin off-resonance frequency and the spherical harmonic components of the eddy currents, and finally also include the *g*-ratio parameters (average log *g*-ratio and *f*_myelin;*k*_).
  e. Steps c-d are repeated until the global minimum is found
4. Finally, we include a fit including all free parameters (both phase- and amplitude-related) initialised from the values found above.

All fits were carried out in python using local optimisation with the quasi-Newton method L-BFGS-B^58,59^ from the scipy library with gradients computed symbolically using the sympy library.

### S3: Unwrapping the phase on a sphere

Another way to avoid the local minima when fitting phase data as discussed in S2 is to unwrap the phase of the data before fitting. Note that as opposed to the more commonly spatial phase unwrapping across an image^55^, we apply phase unwrapping here across the gradient orientations within each voxel.

Similarly, to the estimation of the *l* = 1 components in S2 we start by defining neighbouring gradient orientations as those connected in a convex hull^57^. Starting from some random gradient orientations, any phase wraps in the neighbouring gradients are corrected by subtracting or adding 2*π* to their phases. For each of the neighbours the algorithm is then repeated and so on, until all the phases for all gradient orientations have been unwrapped.

This approach is only expected to work if the SNR is consistently high enough to produce reliable phase estimates for all gradient orientations. This is the case for the phantom data, where we apply phase unwrapping, however it will not generally be the case for DIPPI data, which is why we do not propose to use phase unwrapping when fitting the DIPPI model (where instead we fit directly to the complex data). This phase unwrapping algorithm could probably be made more accurate by adopting some of the techniques used in Jezzard and Balaban (1995) ^55^, but we do not explore that here.

